# The 5’-3’ exoribonuclease XRN4 modulates the plant circadian network in Arabidopsis

**DOI:** 10.1101/2022.07.06.499002

**Authors:** Daniel A. Careno, Soledad Perez Santangelo, Richard C. Macknight, Marcelo J. Yanovsky

## Abstract

Circadian rhythms enable organisms to anticipate and adjust their physiology to periodic environmental changes. These rhythms are controlled by biological clocks that consist of a set of clock genes that regulate each other expression. Circadian oscillations in mRNA levels require regulation of mRNA production and degradation. While transcription factors controlling clock function have been well characterized from cyanobacteria to humans, the role of factors controlling mRNA decay is largely unknown. Here, we show that mutations in XRN4, the central component of the 5’-3’ mRNA decay pathway, alter clock function in Arabidopsis. We found that xrn4 mutants display long period phenotypes for clock gene expression and for the rhythm of leaf movement. These circadian defects were associated with changes in the circadian phases, but not overall mRNA levels, of several core clock genes. We then used non-invasive transcriptome-wide mRNA stability analysis to identify genes and pathways regulated by XRN4. Among genes affected in the xrn4 mutant at the transcriptional and post-transcriptional level, we found an enrichment in genes involved in auxin, ethylene, ABA signaling, and also circadian rhythmicity, although no significant effects were observed for canonical core-clock genes. Strikingly, the mRNAs of several clock regulated BBX genes were stabilized in xrn4 mutants. Some of these BBX genes are auxiliary factors controlling the pace of the clock and are candidates to mediate XRN4 effects on circadian period. Our results establish that, in Arabidopsis, the control of 5’-3’ mRNA decay by XRN4 constitutes a novel post-transcriptional regulatory layer of the circadian gene network.

## Introduction

The circadian clock is an endogenous time-keeping mechanism found in organisms as diverse as bacteria, plants, and animals (Young and Kay, 2001). The clock generates biological oscillations known as circadian rhythms that regulate many aspects of an organism’s behaviour, physiology and development. The clock helps organisms to cope with daily changes in the environment, increasing their fitness (Dodd et al., 2005; Yerushalmi and Green, 2009). In Arabidopsis plants, the circadian clock regulates more than 30% of gene expression (Romanowski et al., 2020). Genes regulated by the circadian clock are essential to many important physiological processes, including, growth, flowering, metabolic regulation and stress responses (Barak et al., 2000).

The Arabidopsis central oscillator is a gene network based on multiple interconnected feedback loops, in which a set of core clock genes acting as transcriptional repressors and activators regulate each other expression, as well as the expression of thousands of clock-controlled genes (Nakamichi, 2020). At dawn, LATE ELONGATED HYPOCOTYL (LHY) and CIRCADIAN CLOCK ASSOCIATED 1 (CCA1)—two MYB domain-containing transcription factors— reach their peak of expression and repress their transcription as well as that of the afternoon and evening genes comprised in the *PSEUDO-RESPONSE REGULATOR* family *(PRR)* and the Evening Complex (EC) (Nagel et al., 2015). Later in the morning, the coactivators NIGHT LIGHT-INDUCIBLE AND CLOCK-REGULATED GENE 1 (LNK1) and LNK2 interact with the single MYB transcriptional activators REVEILLE 8 (RVE8), RVE6, and RVE4 (Xie et al., 2014; Pérez-García et al., 2015), and promote the expression of *PRR5, TIMING OF CAB EXPRESSION 1* (*TOC1*, also known as *PRR1*) and the EC member *EARLY FLOWERING 4* (*ELF4*). Throughout the day the members of the PRR family, starting with PRR9 and ending with TOC1, reach their respective peak of expression and sequentially suppress the expression of *LHY* and *CCA1*. At night, ELF4, ELF3 and LUX ARRHYTHMO (LUX) associate forming the EC. The EC maintains the repression of the *CCA1*, *LHY*, *PRR9*, *PRR5* and *LUX* itself. At the end of the night, CCA1 and LHY repress the EC and the cycle starts again.

Increasing evidence indicates that, besides the transcriptional control of the clock gene network, post-transcriptional mechanisms also play a key role in modulating the function of core clock genes (Beckwith and Yanovsky, 2014). However, while we know in detail the factors regulating the circadian network at the transcriptional level, we are only starting to identify and characterize the factors and pathways controlling this network at the post-transcriptional level.

Alternative splicing (AS), alternative polyadenylation, nonsense-mediated decay (NMD), N^6^ methyladenosine (m^6^A) modification, and nuclear export of mRNAs are among the post-transcriptional mechanisms that regulate circadian clock gene expression (Beckwith and Yanovsky, 2014). For example, *TOC1*, *ELF3*, *CCA1*, *LHY*, *PRR7* and *RVE8* are some of the clock genes undergoing temperature regulated AS. NMD controls the levels of *TOC1* and *ELF3* mRNA (James et al., 2012; Kwon et al., 2014), and light perceived by cryptochromes controls m^6^A deposition in CCA1 mRNA (Wang et al., 2021). Several mutants involved in the nuclear exportation of mRNA are defective in clock function (Ziemienowicz et al., 2003; MacGregor et al., 2013; de Leone et al., 2020). In addition, splicing factors or modulators that impact clock gene expression and function have also been characterized. The first splicing modulator affecting clock function that was identified was PROTEIN ARGININE METHYLTRANSFERASE 5 (PRMT5), whose methylation targets include core spliceosomal proteins, and its activity is necessary for the correct splicing of *PRR9* mRNA (Deng et al., 2010; Sanchez et al., 2010). Among the PRMT5 splicing-related targets, one Sm-like protein (LSM) stands out, LSM4. This protein is required for the formation of two heptameric complexes. On one hand, it is a member of the LSM2-8 nuclear complex that is part of the U6 small nuclear ribonucleoprotein particle, which is the catalytic centre of the spliceosome (Perea-Resa et al., 2012). On the other hand, LSM4 is part of the LSM1-7 cytoplasmic complex involved in mRNA decapping followed by 5’-3’ mRNA degradation (Montemayor et al., 2020). Mutations in *LSM4* and *LSM5* lengthen the period of circadian rhythms in Arabidopsis (Perez-Santangelo et al., 2014). These phenotypes were linked to defective splicing of a sub-set of core clock genes. However, given that LSM4 and LSM5 are part of both LSM2-8 and LSM1-7 complexes, their effect on clock function might also be associated, at least in part, with their role in mRNA decay.

High amplitude mRNA oscillations, such as those observed for core clock genes and several clock-controlled output genes, require regulation of both mRNA stability and mRNA synthesis. Indeed, in a study seeking to identify unstable mRNAs using microarrays, Gutierrez et al. showed that genes controlled by the circadian clock are particularly unstable (Gutierrez et al. 2002). Also, the clock-controlled CCR-LIKE (CCL) and SENESCENCE ASSOCIATED GENE 1 transcripts are differentially regulated at the level of mRNA stability at different times of day (Lidder et al., 2005). Furthermore, light modulates CCA1 mRNA stability, and this contributes to clock entrainment (Yakir et al., 2007, Wang et al., 2021).

Two main mRNA decay pathways shape transcript stability. Both of these pathways begin with the conversion of polyadenylated to oligoadenylate mRNA, through deadenylation or shortening of the polyA tail. Then, mRNAs can be the substrate of the 3’-5’ decay pathway and be degraded by the highly conserved multi-subunit complex called exosome. Alternatively, the 5’ cap can be removed and decapped transcripts can be degraded by 5’-3’ exoribonucleases (XRNs) (reviewed in Zhang and Guo, 2017). In Neurospora, a role for the catalytic component of the exosome complex in the control of circadian rhythms was described several years ago (Guo et al., 2009). Whether the 5’-3’ mRNA decay pathway controls clock function in eukaryotic organisms has yet to be shown.

The Arabidopsis genome encodes three XRNs: the nuclear XRN2 and XRN3, and the cytoplasmic XRN4, which is the functional homolog of yeast and mammalian XRN1 and ortholog of the nuclear XRN2/RAT1 (Kastenmayer and Green, 2000). XRN4 main role is mRNA degradation, which must be preceded by decapping or endonucleolytic cleavage. Mutants for *XRN4* were independently isolated in two screening for mutants insensitive to ethylene (Van Der Straeten et al., 1993; Ecker, 1995), which were later associated with the 5’-3’ exoribonucleases 4 gene (*XRN4*) (Olmedo et al., 2006; Potuschak et al., 2006).

Mutants for *XRN4* show accumulation of deadenylated uncapped transcripts and 3’ mRNA fragments that are subject to nonsense-mediated decay (NMD) (Nagarajan et al., 2019). It was also found that polyA+ mRNA targets of XRN4 are subject to co-translational decay, and this modulates their translation efficiency during plant development (Carpentier et al., 2020). Although targets for XRN4 have been discovered, the overall overlap of target genes between the different studies is low (Rymarquis et al., 2011; Nagarajan et al., 2019; Carpentier et al.,2020). This low overlap could be due to differences in XRN4 activity at different developmental stages.

Here we show that mutations in *LSM1A1B*, which affect decapping, and in *XRN4*, which impacts 5’-3’ mRNA degradation, lengthen circadian period in Arabidopsis. We also observed changes in the circadian phases rather than in the overall mRNA levels of canonical core clock genes in *xrn4* mutants compared to WT plants. Finally, by performing a non-invasive genome-wide analysis of post-transcriptional regulation in *xrn4* mutants and WT plants, we found an enrichment in the circadian rhythm category among those genes with evidence of altered stability in *xrn4* mutants, suggesting a role for 5’-3’exoribonucleases in the control of mRNA stability of some clock regulated transcripts. Indeed, several members of the *BBX* gene family that display high amplitude circadian oscillations and control clock function showed enhanced mRNA stability in *xrn4* mutants and could mediate in part XRN4 effects on the clock. Although the exact and precise molecular mechanisms through which mRNA decay affects the biological clock remain elusive, our results support an important role for the 5’-3’ mRNA decay pathway controlled by XRN4 in the fine-tuning of circadian oscillations in Arabidopsis plants.

## Results

### Lack of Arabidopsis mRNA decay components LSM1 and XRN4 alter circadian rhythms

LSM1 is the defining component of the LSM1-7 complex involved in 5’-3’ mRNA decay and, in Arabidopsis, is encoded by two genes, *LSM1a* and *LSM1b*. Circadian phenotypes were not observed in either single *lsm1a* or *lsm1b* mutants, or in homozygous *lsm1a*/heterozygous *lsm1b* mutant plants (Supplemental Figure S1). Homozygous *lsm1a*/*lsm1b* (*lsm1a/b*) mutants, like *lsm4* (Perez-Santangelo et al., 2014), display severe developmental retardation compared to wild-type (WT) plants. Therefore, circadian rhythms in *lsm1a/b* mutants were evaluated using bioluminescent reporters rather than leaf movement assays. For this, we transformed plants that were homozygous for *lsm1a* and heterozygotes for *lsm1b* with two different circadian bioluminescent reporter genes, pCCA1:LUC and pCCR2:LUC, and analyzed circadian rhythms in homozygous *lsm1a lsm1b* plants in the next generation. The circadian period of bioluminescence was lengthened in *lsm1a1b* homozygous mutants relative to WT plants for both pCCA1:LUC (25.11 ± 0.40 h and 24.50 ± 0.44 h, respectively, p<0.001) and pCCR2:LUC reporters (25.70 ± 0.48 h and 24.62 ± 0.33 h, respectively; p<0.001), suggesting that the 5’-3’mRNA decay pathway controls clock function in Arabidopsis (Figure 1). We then evaluated if *LSM1a* and *LSM1b* mRNA levels were under the control of the circadian clock, using publicly available datasets. We found that *LSM1b* displays circadian oscillations in the DIURNAL dataset (Mockler et al., 2007), but none of the *LSM1* genes cycle in our recently published RNAseq circadian time-course (Romanowski et al., 2020), suggesting weak control of *LSM1* expression by the circadian oscillator.

**Figure 1.**
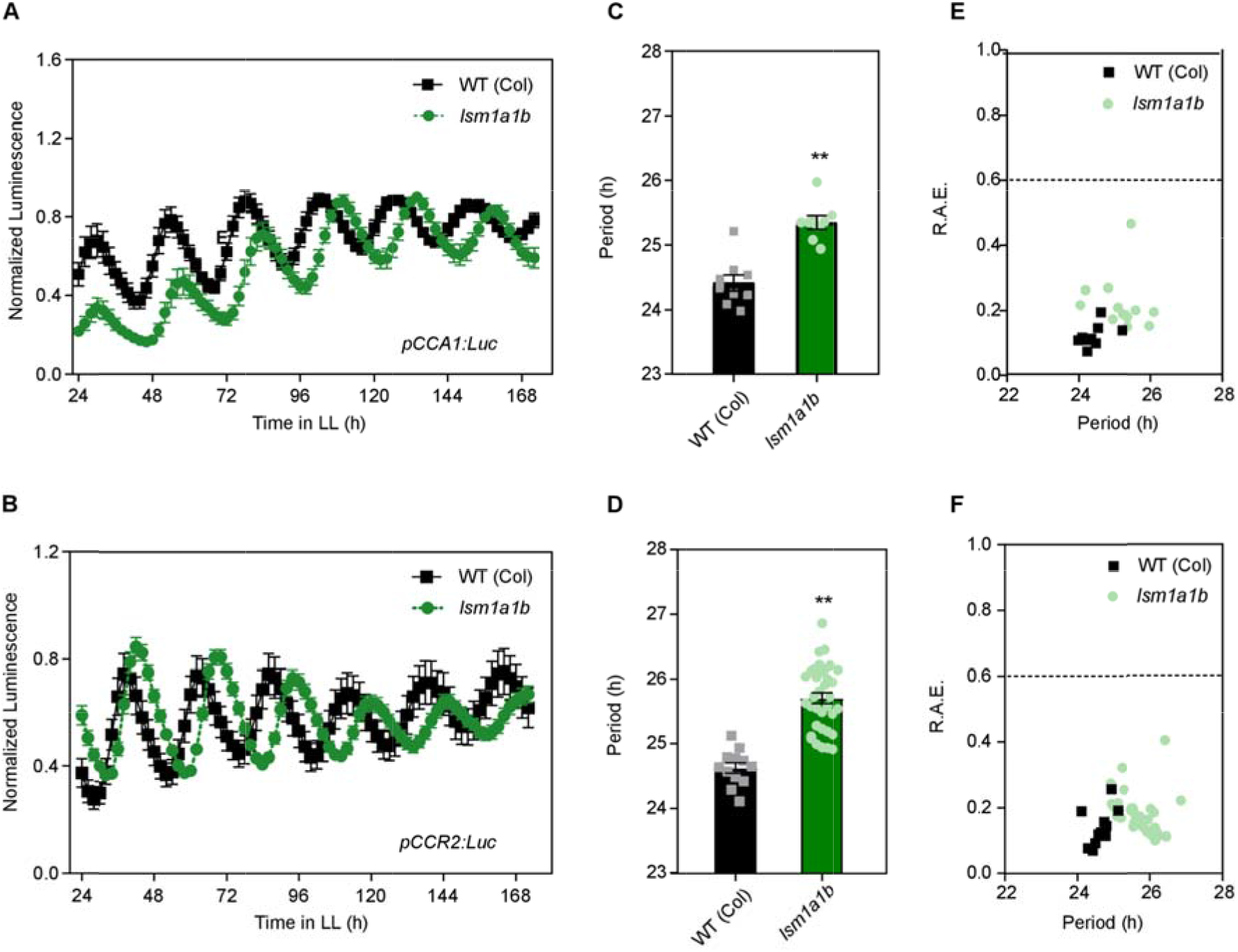
Mutations in the plant *LSM1a/LSM1b* genes affect the clock. The expression of the morning core clock reporter pCCA1:LUC (A) and afternoon clock output reporter pCCR2:LUC (B) was measured for 7 d in constant light (LL) after entrainment under long-day (LD) conditions. WT (Col), wild-type Columbia ecotype (black squares); *lsm1a/b* mutant (green circles). Bioluminescence was recorded every 2 h over the 7 days and analysed by FFT-NLLS. (C,D) Period values and (E,F) Relative Amplitude Error (R.A.E) were plotted for each genotype. Error bars represent themean with range. **p < 0.001 (Student t test).

After removal of the 5’ 7-mG cap by the decapping complex, the unprotected 5’ end of mRNAs can be degraded by specific 5’-3’exoribonucleases. In Arabidopsis, XRN4 is the only 5’-3’exoribonuclease with a cytoplasmic location. To analyze the effect of cytoplasmic mRNA decay on clock function, we characterized circadian rhythms in *xrn4* single mutants. Using a bioluminescent reporter, we found that the circadian period of *TOC1* expression was lengthened in two different *xrn4* mutants relative to WT plants (25.59 ± 0.52h for *xrn4-3* compared to 24.14 ± 0.42h for WT plants, p <0.001, Figure 2 A, B, C; and 25.87 ± 0.08 h for *xrn4-5* vs 25.17 ± 0.26 h for the WT, p<0.05, Supplemental Figure S2). We also observed a longer circadian period for the rhythm of leaf movement in the two different *xrn4* mutants compared to WT plants (25.27 ± 0.33h for *xrn4-3*, 24.76 ± 0.29 h for *xrn4-5* and 24.00 ± 0.26 h, for WT plants; p < 0.001) (Figure 2 D, E, F). Altogether these results provide compelling evidence that the canonical mRNA 5’-3’ decay pathway, which includes the LSM1-7 complex and the 5’-3’exoribonuclease XRN4, plays an important role in the control of circadian period in Arabidopsis plants.

**Figure 2.**
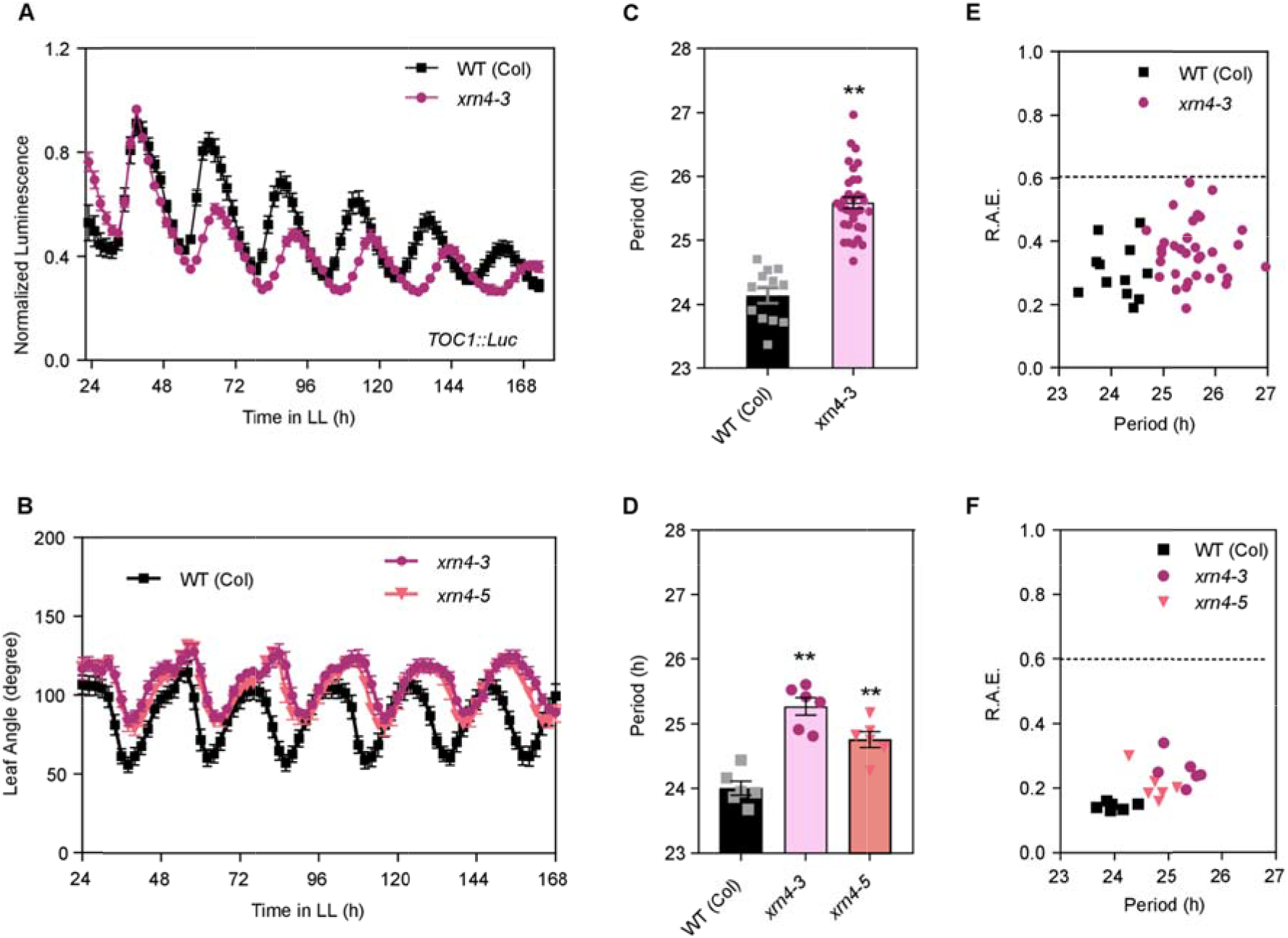
Mutations in the exoribonuclease 5’-3’ *XRN4* cause a delay in clock period. (A) Expression of the night core clockreporter pTOC1:LUC was measured for 7 d in constant light (LL) after entrainment under LD conditions. WT (Col), wild-type Columbia ecotype(black squares); *xrn4-3* mutant (purple circles). (B) Circadian rhythm of leaf movement. Leaf angles weremeasured for the first pair of leaves in seedlings entrained under long-day (LD) conditions and plants were then transferred to constant light (LL). WT (Col) (black squares). *xrn4-3* mutant (purple circles, n = 6); *xrn4-5* mutant (red triangle). (C,D) Period length of bioluminescence and leaf movement rhythms wasestimatedby fast Fourier transform nonlinear least test (FFT-NLLS). (E,F) Relative Amplitude Error (R.A.E) were plotted for each genotype. Error bars represent the mean with range. **p < 0.001 (Student t test).

### Loss of Arabidopsis *XRN4* alters the timing of expression of core clock genes

To further study the effect of the mRNA decay pathway on clock function, and since XRN4 is the only cytoplasmic enzyme involved in the 5’-3’ mRNA decay, we decided to analyse the expression of several core clock genes in the *xrn4-3* mutant. For this, plants were grown under light/dark cycles for 10 days and then transferred to constant light (LL) for 3 d. Samples were collected every 4h for 1d. Phase changes were observed for most of the core clock genes evaluated in *xrn4-3* mutant and WT plants. For the morning genes *LHY* and *CCA1*, a reduced and delayed expression was observed in *xrn4-3* mutants compared to WT plants, resulting in loss of the sharp morning peak (Figure 3 A, B; p<0.05). In the case of the midday/evening genes *PRR9, PPR5* and *ELF3* a clear 4h phase delay was also observed in *xrn4-3* compared to WT plants (Figure 3 C, E and H), resulting in a higher and broader peak of expression for *PRR5* in *xrn4-3* than in WT plants. For *TOC1* no differences in phase of expression were found between genotypes, but a higher peak of expression was detected in *xrn4-3* (Figure 4 F). For the morning gene *RVE8*, we observed a slightly higher mRNA levels in the *xrn4* mutant compared with WT plants during subjective daytime (Figure 3 G; p<0.05). Last, no differences in *PRR7* expression were observed between *xrn4-3* mutants and WT plants under these conditions (Figure 3 D). In summary, most genes showed loss of a sharp peak of expression, resulting in a wider peak of expression during the third day under constant conditions, which is compatible with the approximately 1-hour period difference observed between *xrn4-3* mutants and WT plants. Strong differences in overall mRNA levels were not observed for most clock genes between genotypes, although a slight enhancement in mRNA levels was observed for *PRR5* and *RVE8* in the decaying phase of expression. These results, taken together, suggest that XRN4 effect on the clock could be in part the result of subtle effects on mRNA levels of several clock genes, rather than the result of a strong effect on the mRNA levels of a few target genes.

**Figure 3.**
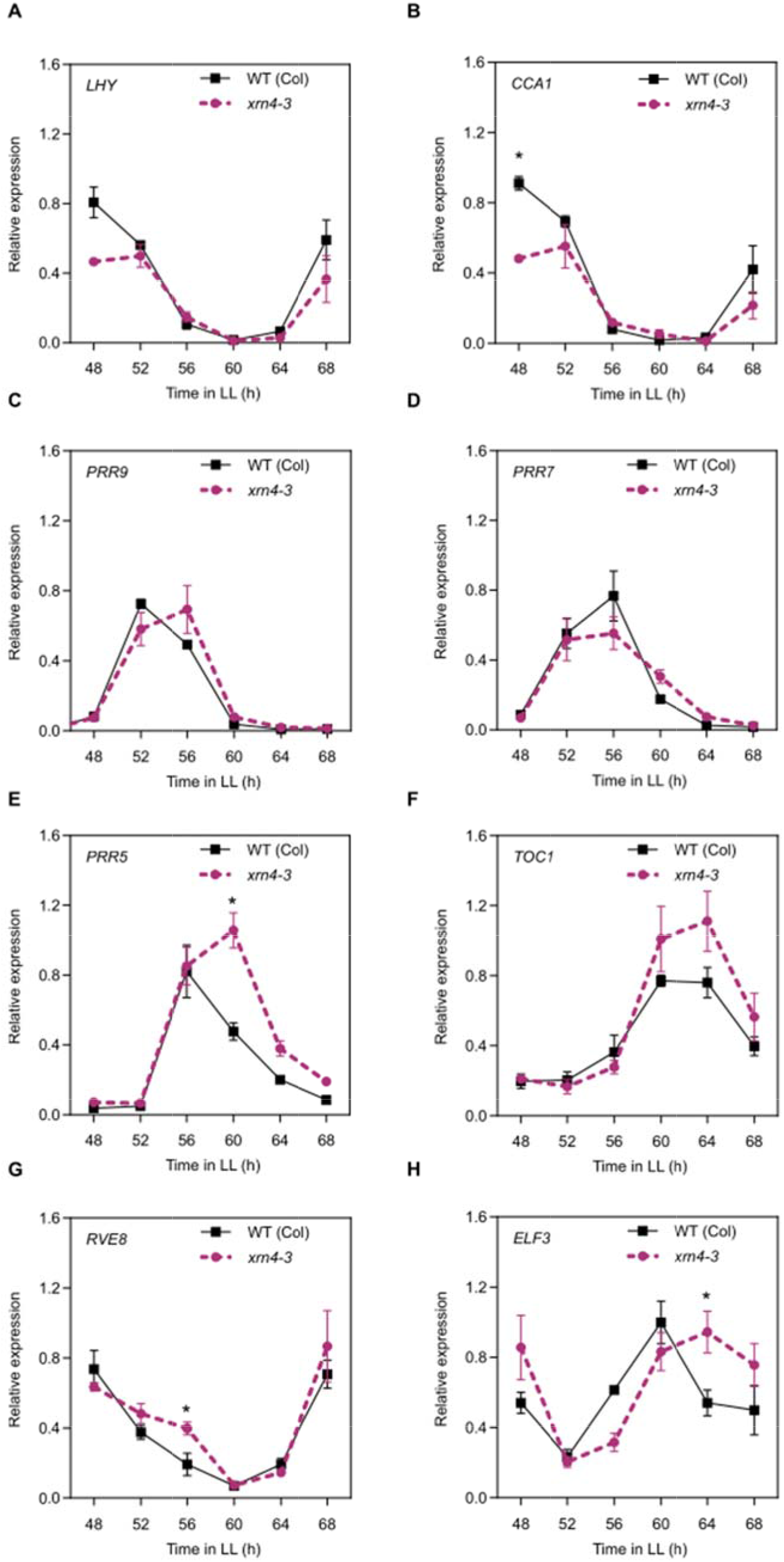
Mutations in the exoribonuclease 5’-3’ *XRN4* affect core clock expression patterns. Gene expression of core clock genes measured by RT-qPCR in plants entrained under 12 h light/12 h darkness, before being moved to constant light (LL) for 3 days. WT (Col) (solid black squares), *xrn4-3* mutant (dashed purple line). (A) LHY, (B) CCA1, (C) PRR9, (D) PRR7, (E) PRR5, (F) TOC1, (G) RVE8, (H) ELF3. Data are the average of three biological replicates normalized to the maximum value. Error bars represent SEM. * p < 0.05 (Student t test).

**Figure 4.**
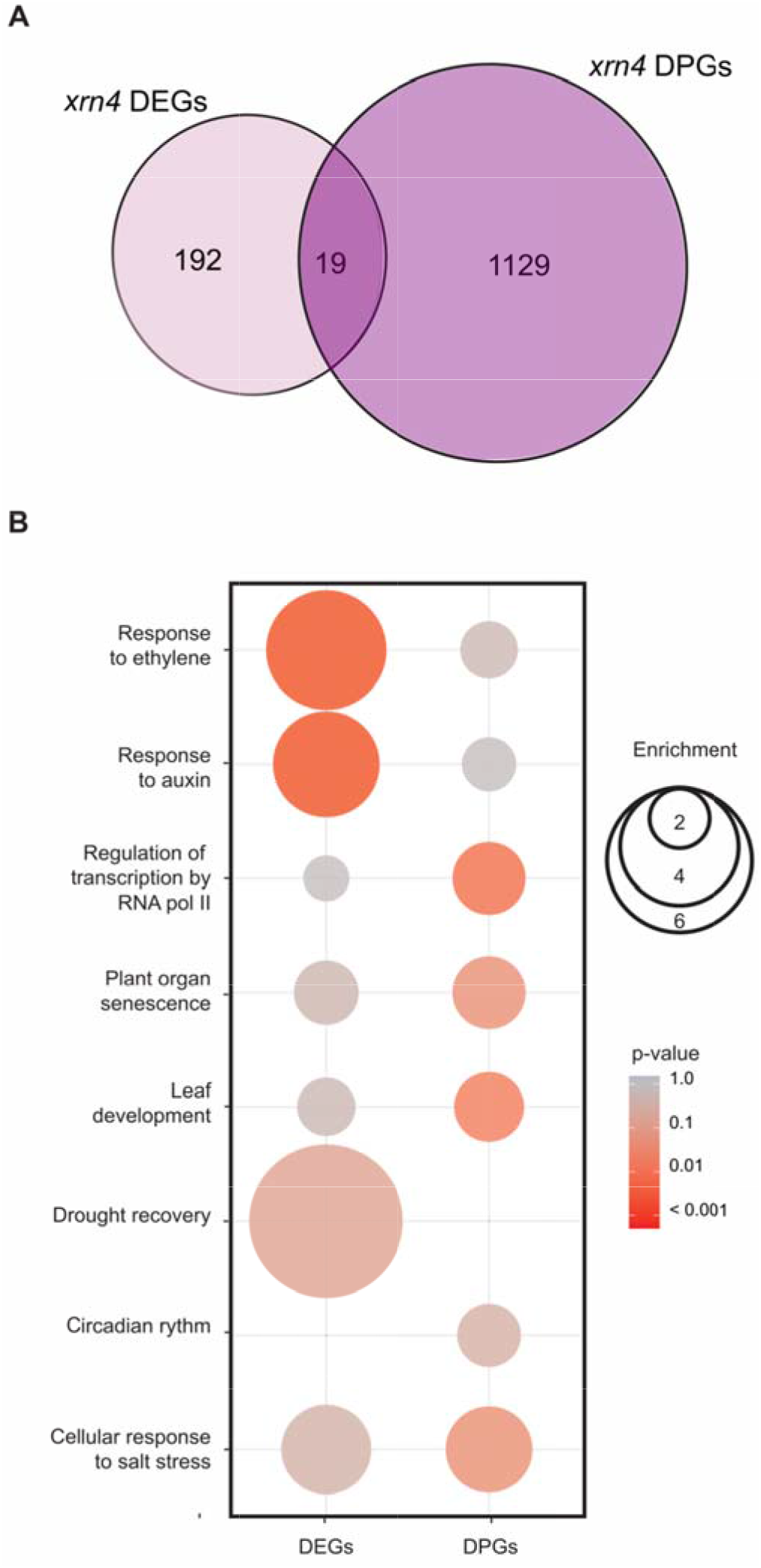
Global analysis of differentially regulated genes at the transcriptional and post-transcriptional level in the 5’-3 ‘exoribonuclease *xrn4-3* mutant. (A) Overlap between differentially expressed genes (DEGs) and differentially post-transcriptional regulated genes (DPGs) between WT (Col-0) and *xrn4-3*. (B) Enrichmentfactor and p-value of selected gene ontologiesamong DEGs and DPGs. The colour gradient represents adjusted P-values and the differences in bubble sizecorrelate with the enrichment factor.

### Flowering time is affected in *xrn4* mutant

To study if flowering time, an output of the circadian clock, was affected by mutations in the cytoplasmic 5’-3’ exoribonuclease, *xrn4* mutant plants were grown under inductive photoperiodic conditions (long days, 16h light and 8h darkness). We observed a late flowering phenotype for *xrn4*, evaluated as the number of rosette leaves at bolting. The *xrn4-3* mutant allele flowered with 22.58 ± 1.03 leaves and the *xrn4-5* allele with 20.0 ± 0.74 leaves, both compared to 15.83 ± 0.56 leaves in WT plants (Supplemental Figure S3; p<0.05). The late flowering phenotype observed for these mutants under long day conditions is consistent with this defect being, at least in part, the result of the longer circadian period of the mutants, although some partial clock independent effects cannot be excluded.

### Genome-wide analysis of transcriptional and post-transcriptional regulation by XRN4

Differences in circadian phase have been shown to strongly affect the analysis of differential gene expression between WT and clock mutant plants, even in mutants with mild circadian period alterations (Hsu and Harmer, 2012). Our finding that *xrn4* mutants display an approximately 1 hour longer circadian period phenotype compared to WT plants, suggests that many of the mRNA previously identified as differentially accumulated in *xrn4* mutants could have simply been selected because of differences in their phase of expression, rather than differences in their overall mRNA levels. Here, we performed a transcriptomic analysis on total RNA using a protocol designed specifically to explore the effects of *xrn4* mutation on overall mRNA levels and minimize a possible impact of period length differences between genotypes. For this, we grew plants for 15 days under long day conditions and transferred them to constant continuous light conditions. Then we harvested samples every 4h, from Circadian Time 2 (CT2, i.e. 2 hours after lights on under continuous light) to CT22. Finally, to conduct a comparative transcriptomic analysis, we pooled equal amounts of samples obtained at the different times of day into one sample for each biological replicate for each genotype. The pooling of samples harvested at different times of day during a circadian cycle should attenuate differences in mRNA levels caused by possible changes in the phase of expression of clock-controlled genes between genotypes, and, therefore, should allow the identification of genes showing different overall mRNA levels between *xrn4* mutant and WT plants.

The analysis of the transcriptomic data revealed 211 differentially expressed genes (DEGs) between WT and *xrn4* mutants (Figure 4A; Supplemental Dataset S1). The majority of these genes, 80% (167/211), were upregulated in the mutant compared to WT plants, as expected for possible targets of the 5’-3’ mRNA decay pathway (Supplemental Dataset S1) (Nagarajan et al., 2019; Carpentier et al., 2020). Interestingly, the group of genes upregulated in *xrn4* mutants showed a 2-fold statistically significant enrichment in high amplitude Circadian Controlled Genes (CCGs; top 500 circadian regulated genes sorted by amplitude, Romanowski et al., 2020) (Supplemental Figure S4 A; Supplemental Table S1). As expected, gene ontology (GO) analysis of DEGs also showed enrichment in terms that correspond to previously reported XRN regulated processes (Figure 4B; Supplemental Dataset S3). This includes responses to the hormones ethylene, auxin and ABA, responses to drought, the regulation of germination, root development and anthocyanin biosynthesis (Van Der Straeten et al., 1993; Ecker, 1995; Windels and Bucher, 2018; Wawer et al., 2018; Basbouss-Serhal et al., 2017; Hirsch et al., 2011 Kwon et al., 2011).

Next, to investigate the impact of *xrn4* mutations on mRNA stability, we performed a non-invasive analysis of differential post-transcriptional regulation using the R package INSPEcT (Furlan et al., 2020). Briefly, INSPEcT detects changes in post-transcriptional regulation by comparing the relationship between intronic and exonic reads, which are used as proxies for pre-mRNA and mature mRNA. Thus, differences in mRNA degradation rates between genotypes can be inferred from changes in the ratio between premature and mature RNA for each gene. Using this approach, we found 1148 differentially post-transcriptional regulated genes (DPGs) in *xrn4* compared to WT plants (Figure 4A; Supplemental Dataset S2)

No canonical core-clock genes were identified among the 1148 genes differentially regulated at the post-transcriptional level for the *xrn4* mutants. An enrichment in the GO term circadian rhythms and other GO terms related to phenotypes previously reported for *xrn* mutants was observed, similar to what we found in the DEG analysis (Figure 4B; Supplemental Dataset S3B). The minor and subtle effects observed at the post-transcriptional level on core clock genes in *xrn4* mutant compared to WT plants using INSPEcT are consistent with the rather similar mRNA decay rates observed for selected core-clock genes in *xrn4* mutants and WT plants after addition of the transcriptional inhibitor cordycepin (Supplemental Figure S5). This suggests that XRN4 might control clock function, in part, through very subtle effects on mRNA decay rates of some core-clock genes. Alternatively, or in addition, XRN4 could be controlling clock function through post-transcriptional regulation of auxiliary clock factors rather than canonical core clock components. Indeed, among the DPGs identified in *xrn4* mutants we found several B-BOX domain-containing genes, *BBX2* (*COL1*), *BBX3* (*COL2*), *BBX11*, *BBX15* and *BBX19*. These genes display high amplitude circadian rhythms in mRNA levels and some of them are known to control circadian period length (Ledger et al., 2001). We analysed the mRNA decay rates of some of these candidate genes in WT and *xrn4* plants, with the cordycepin-induced transcription arrest assay, and observed a tendency towards slower mRNA decay rates for *BBX3* and *BBX15* genes in the mutant plants (Figure 5).

**Figure 5.**
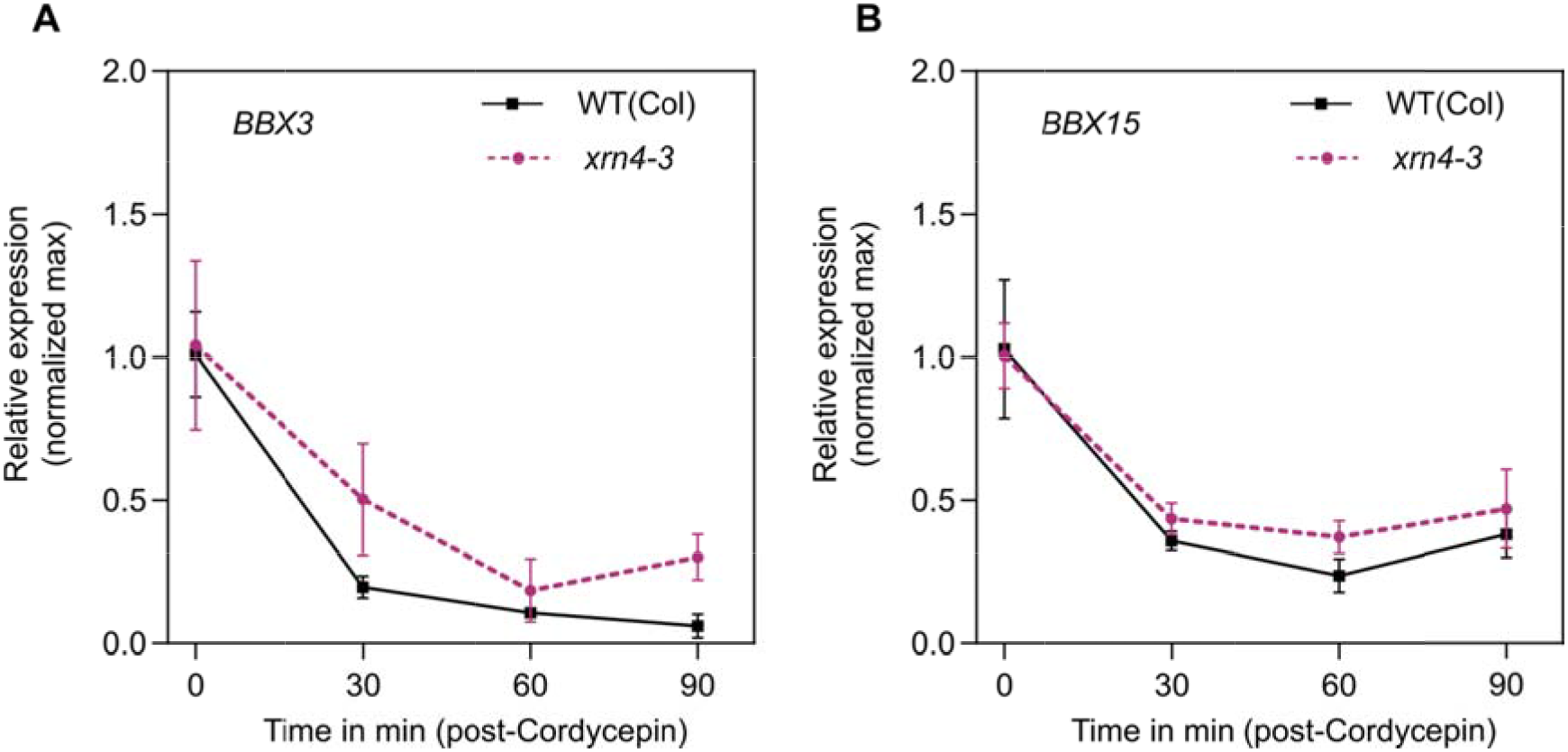
mRNA decay rates for the candidates BBX3 and BBX15 in *xrn4-3*. Analysis of mRNA decay with a cordycepin-induced transcription arrest assay. Plant where grew for 10 days in continuous light and samples were taken every 30min from 0 to 90 min. (A) BBX3, (B) BBX15.

An examination of the ratio between mature and premature RNA levels (M/P) in genes that were characterized as DPGs in *xrn4* mutant plants revealed that 90% of the genes (972 out of 1148) exhibited greater M/P ratios in *xrn4* mutants than in WT plants (Supplemental Figure S6). This is the expected behaviour for true targets of the XRN4 exoribonuclease, since reduced degradation of an XRN4 target in *xrn4* mutants should result in higher M/P ratios. This trend extends to the DPGs included in the GO term circadian rhythms (Supplemental Figure S7). Altogether, these results support the conclusion that mRNA-decay is an important post-transcriptional regulatory layer of the Arabidopsis circadian network.

## Discussion

Although the importance of post-transcriptional regulation in shaping circadian oscillations of gene expression is well recognised, the role of mRNA decay has been largely unexplored (Beckwith and Yanovsky, 2014; Lück et al., 2014; Romanowski and Yanovsky, 2015; Mateos et al., 2018). So far, only one study has revealed a connection between the 3’-5’mRNA decay machinery and the control of circadian rhythms in Neurospora (Guo et al., 2009). Here we show for the first time, evidence that the 5’-3’ mRNA decay pathway is also involved in fine-tuning circadian rhythms.

We have previously described that two components of the U6 snRNPs spliceosomal complex, LSM4 and LSM5, affected clock function in Arabidopsis, most likely by regulating the alternative splicing of a sub-set of core clock genes (Perez-Santangelo et al., 2014). Since LSM4 and LSM5 are not only involved in nuclear RNA processing, but also modulate mRNA decapping in the cytoplasm, we asked whether the circadian phenotypes could be due, in part, to effects of LSM4 and LSM5 on mRNA decay. To address this, we took advantage of the fact that LSM1 is the defining component of the cytoplasmic LSM1-LSM7 decapping complex that modulates mRNA decay and it is not present in the LSM2-LSM8 nuclear spliceosome complex. We found that mutations in the Arabidopsis *LSM1a/LSM1b* genes resulted in a long circadian period phenotype (Figure 1). In human cells, ribosome profiling revealed that the LSM1 protein is rhythmically translated and that cytoplasmic P-body formation was rhythmic in these cells (Jang et al. 2015), which could be contributing rhythmicity to the 5’-3’ mRNA decay process.

After removal of the 5’ 7-mG cap by the decapping machinery, mRNAs are degraded in the cytoplasm by XRN exoribonucleases. A triple mutant for the three Arabidopsis *XRN* genes, *xrn2/3/4*, has been shown to have a long period phenotype for the expression of several core clock genes (Litthauer et al., 2018). However, which of these XRNs contributes to the clock phenotype was not known. XRN4 is the only 5’-3’exoribonuclease located in the cytoplasm where mature mRNAs are degraded, whereas XRN2 and XRN3 contribute to the processing and degradation of several classes of nuclear RNAs. Supporting a role for the canonical cytoplasmic mRNA decay pathway in the control of clock function, we found that mutations in *XRN4* cause an approximately 1h delay in the circadian period of the rhythm of leaf movement and *TOC1* promoter activity (Figure 2). We also observed a reduction or shift in the expression profiles of the majority of core clock genes analysed under free running conditions (Figure 3). Particularly for *ELF3* and *PRR5*, the approximately 4-hour phase delay and widening of the peak of expression observed on the third day under continuous light, is consistent with the approximately 1 h increase in period length present in *xrn4* mutants and could be interpreted as the result of slower mRNA decay of clock genes.

Several genome-wide analyses of XRN4 effects on the Arabidopsis transcriptome have been conducted before (Rymarquis et al., 2011, Nagarajan et al., 2019, Carpentier et al., 2020). The transcriptomic analysis of mutant plants with circadian period phenotypes requires special care to avoid identifying genes affected in their circadian phases of expression, rather than in their overall mRNA levels, which happens when samples collected at a singly time point during a daily cycle are evaluated (Hsu and Harmer, 2012). This prompted us to conduct a new genome wide analysis to study overall changes in mRNA levels in the Arabidopsis *xrn4* mutants by comparing mRNAs from samples collected throughout a full circadian cycle, rather than at a single time point. This approach revealed 167 genes whose overall mRNA levels were higher in *xrn4* mutants than in WT plants. Several of these genes were previously reported to be regulated by XRN4, as expected. We observed an enrichment in genes associated with auxin and ethylene signalling, pathways already known to be modulated by XRN4. Indeed, a role for XRN4 (*ein5*, ethylene insensitive 5) in the regulation of ethylene responses is well known (Olmedo et al., 2006; Potuschak et al., 2006). Furthermore, ethylene emissions display circadian oscillations (Thain, 2004) and the *ein3* mutant, which is also involved in the ethylene pathway, presents a long period phenotype (Haydon et al., 2017). Therefore, it is possible that at least part of the *xrn4* circadian phenotypes could be associated with the role of XRN4 in modulating the ethylene pathway. It is important to mention that the precise mechanisms through which ethylene signalling controls the clock is not yet known.

The differential expression gene analysis (DEG) on overall mature mRNAs showed no core-clock genes among those upregulated in *xrn4* mutants, suggesting that XRN4 effects on mRNA levels of core-clock genes are subtle, as observed in our qRT-PCR timecourse analysis. Interestingly, we did find a similar and significant enrichment of genes displaying high amplitude circadian oscillations in mRNA levels among those upregulated in *xrn4* mutant (Supplemental Figure S7; Table S3). Finally, using the same RNA-seq dataset generated here, we performed an analysis of XRN4 effects on post-transcriptional regulation. For this, we used the Bioconductor package INSPEcT, a non-invasive approach that allows the detection of changes in transcription and decay rates of mRNA levels, by comparing the relationship between intronic and exonic reads from RNA-seq data generated using Total RNA (Furlan et al., 2020). This analysis showed 1148 post-transcriptionally regulated genes (DPGs) for *xrn4* (Figure 4B, Supplemental Figure S7). Noteworthy, we found more DPGs than DEGs in our RNA-seq dataset, consistent with a predominant role for XRN4 in post-transcriptional regulation. Many DPGs were not identified as DEGs (Figure 4; Supplemental Dataset S1 and S2). This might be due to a buffering system involving interactions between post-transcriptional and transcriptional regulatory mechanisms. For example, deletion of XRN1 in yeast has been shown to alter transcription rates (Braun and Young, 2014). On the other hand, *xrn4* mutant plants show an increase in Post-transcriptional gene silencing (PTGS; Gazzani et al., 2004).

Among new genes identified as potential targets of XRN4 in our analysis, we focused our attention on the family of BBX transcription factors, due to the appearance of multiple of its members among DEGs and DPGs (Supplemental Dataset S1 and S2). For example, *BBX15, BBX2 (COL1*) and *BBX3 (COL2*) were identified as DPGs in *xrn4* mutant and overexpression of COL1 and COL2 alters circadian period (Ledger et al., 2001). Moreover, we observed a tendency towards slower mRNA decay rate for *BBX3* and *BBX15* after *c*ordycepin-induced arrest of transcription in *xrn4* mutant (Figure 5). Last, *BBX19* was also identified as a DPG in *xrn4*, and has recently been found to interact with the *PRR* gene family forming a transcriptional regulatory complex that acts fine-tuning the circadian clock (Yuan et al., 2021). Whether part of *xrn4* circadian phenotype is due to altered regulation of these *BBX* genes remains to be determined.

In conclusion, circadian regulation of mRNA levels requires regulation of both mRNA synthesis and mRNA degradation, but the specific mRNA decay factors involved in the control of circadian rhythms in multicellular organisms was not known. Here we provide strong evidence that LSM1 and XRN4, components of the 5’-3’mRNA decay pathway, contribute to fine-tuning the Arabidopsis circadian clock. This regulation involves subtle alterations in the abundance of core-clock genes, as well as global effects on overall mRNA accumulation of high amplitude clock-controlled genes (CCGs). Further studies are needed to understand the precise mechanisms through which the 5’-3’ mRNA decay pathway regulates this key biological process.

## Materials and Methods

### Growth Conditions

Plants were grown in pots containing a mixture of organic substrate, vermiculite, and perlite [0.5:1:1 (vol/vol)] at 22 °C under long days (LD; 16-h light/8-h dark cycles), 12:12 (12h light/12hs dark cycles) or continuous light (LL), depending on the experiment.

### Plant Material

The mutants used, including *lsm1a* (SALK_106536), *lsm1b* (SAIL_898_D06), *xrn4-3* (SALK_014209), *xrn4-5* (SAIL_681_E01), were obtained from the Arabidopsis Biological Resource Center (ABRC).

### Circadian Leaf Movement Analysis

For leaf movement analysis, plants were entrained under a 16-h light/8-h dark cycle, transferred to continuous white fluorescent light, and the position of the first pair of leaves was recorded every 2 h for 5–6 d using Image J software (imagej.nih.gov/ij/). Period estimates were calculated using Brass 3.0 software (Biological Rhythms Analysis Software System; www.amillar.org) and analyzed using FFT-NLLS. The statistical analysis was done using a two-tailed Student t test or one-way ANOVA, depending on the case.

### Flowering Time Analysis

For flowering time analysis, plants were grown under long days (16 h light/8 h dark) at a constant temperature of 22 °C. Flowering time was estimated by counting the number of rosette leaves at the time of bolting. The statistical analysis was done using a two-tailed Student t test.

### Plant Bioluminescence Recording and Data Analysis

The seedlings were grown directly on Murashige and Skoog 0.8% agar supplemented with 1% sucrose in a 96-well plate. One seed was placed per well, and the plate was placed in darkness at 4 °C for 3 d. Then seedlings were entrained under long days (16-h light/8-h dark cycle). After 7 d, the entire plate was transferred to constant light and temperature (22 °C) conditions and placed in a luminometer to measure bio-luminescence emitted by each seedling every 2 h. After 5–6 d, analysis of the data was done using the software Mikrowin 2000 (version 4.29) (Mikrotek Laborsysteme). The luminometer model used was the microplate reader Centro LB-960 (Berthold Technologies). Period estimates were calculated with Brass 3.0 software (Biological Rhythms Analysis Software System; www.amillar.org) and analyzed using FFT-NLLS. The statistical analysis was done using a two-tailed Student t test.

### Cordycepin treatment

Plants were grown in continuous light (LL) since germination, with no entrainment. Under LL gene oscillations are flatten and changes in overall expression can be seen no matter the time of the day. After two-week, seedlings were transferred to flasks containing a buffer (1mM PIPES, pH 6.25, 1 mM sodium citrate, 1 mM KCl, 15 mM sucrose), and after a 30-min incubation, the first sample was taken (time =0h). Cordycepin (150 mg/L) was added and the rest of the samples were taken at 30min, 60min and 90min. Total RNA was extracted using Trizol reagent (Invitrogen) for quantitative RT-PCR.

### Quantitative RT-PCR

For time course analysis, 14-d-old plants were grown under 12-h light/12-h dark cycles and then transferred for 3 d to continuous white light at 22 °C. Samples were collected every 4 h for 1 d. Total RNA was obtained from these samples using TRIzol reagent (Invitrogen). One microgram of RNA was treated with RQ1 RNase-Free DNase (Promega) and subjected to retro-transcription with M-MLV (Invitrogen) and oligodT according to the manufacturer’s instructions. Synthesized cDNAs were amplified with FastStart Universal SYBR Green Master (Roche) using the Mx3000P Real Time PCR System (Agilent Technologies) cycler. The PP2A (AT1G13320) transcript was used as a housekeeping gene. Quantitative RT-PCR quantification was conducted using the standard curve method as described in the Methods and Applications Guide from Agilent Technologies. Primers used for amplification of each gene are described in Supplemental Table S2.

### Growth conditions and protocol used for total RNA library preparation and high-throughput sequencing

Four biological replicates of WT (Col-0), *xrn4-3* seeds were sown onto Murashige and Skoog medium containing 0.8% (w/v) agarose, stratified for 3 days in the dark at 4°C, and then grown at 22°C in long day conditions (16h light, 8h dark). After two weeks, plants were transferred to continuous light. During the first day of continuous light whole seedlings were harvested every 4 hours starting at CT=2hs up to CT=22hs. Total RNA was extracted with TRIzol (Invitrogen) following the manufacturer’s protocols. Samples were pooled across the time series before library preparation to estimate the difference in absolute RNA levels among genotypes, rather than differences resulting from alterations in the timing of expression of each gene.

For library preparation, ribosomal RNA was removed by Ribo-Zero™ Magnetic Kit (Illumina) and rRNA free residue was cleaned up by ethanol precipitation. Subsequently, sequencing libraries were generated using Directional NEBNext Ultra RNA Library Prep Kit (NEB) (Novogene, Singapore). Samples sequenced on the Illumina NovaSeq 6000 platform, providing 150bp paired-end reads (Novogene, Singapore). On average 108 million 150-nt paired-end reads were obtained for each sample.

### Processing of RNA sequencing reads

Sequence reads were mapped to Arabidopsis thaliana TAIR10 (Lamesch et al., 2012) genome using STAR 2.7.0 (Dobin et al., 2013) with the following parameters: --twopassMode Basic --outFilterMultimapNmax 2 --outFilterType BySJout --outSJfilterReads Unique --sjdbOverhang 149 --alignSJoverhangMin 6 --alignSJDBoverhangMin 3 -- alignIntronMin 20 --alignIntronMax 5000.

### Differential gene expression analysis

Before differential expression analysis, we discarded genes with fewer than 10 reads on average per condition (minRds>=0.05). Differential gene expression was estimated using the ASpli package version 2.0.0 (Mancini et al., 2021) using custom R scripts, and resulting p-values were adjusted using a false discovery rate (FDR) criterion (Benjamini and Hochberg, 1995). In order to detect the subtles effects of XRN4 we filter genes based on a low stringency FDR and log2 fold change. Genes with FDR values lower than 0.1 and an absolute log2 fold change greater than 0.58 were deemed differentially expressed. Differentially expressed genes (DEGs) in the *xrn4-3* mutant compared to WT (Col-0) can be found in Supplemental Dataset 1.

### Differential gene post transcriptional regulation analysis

Differential post-transcriptional regulated genes (DPGs) were estimated using the R package INSPEcT version 1.22.0 (Furlan et al., 2020). Expression was quantified from BAMs using the built-in function with default parameters. Genes with a log2 normalized expression lower than 0.25 were discarded. DPGs were determined using compareSteadyNoNascent() using a log2 distance threshold of 0.58. DPGs in the *xrn4-3* mutant compared to WT (Col-0) can be found in Supplemental Dataset 2.

### Gene Ontology (GO) analysis

GO terms assignment for the DEGs and DPGs datasets were obtained using the Bioconductors packages topGO version 2.44.0 and org.At.tair.db version 3.13.0. An enrichment test was performed for the biological process category. P-values were obtained using the Fisher Exact Test. The enrichment factor (EF) was estimated as the ratio between the proportions of genes associated with a particular GO category present in the dataset under analysis, relative to the proportion of the number of genes in this category in the whole genome that have passed the expression filters.

### Clock Controlled Genes (CCG) overrepresentation analysis

Similar to the GO analysis, for the DEGs and DPGs datasets we calculated an enrichment factor for clock-controlled genes with a relative amplitude greater than 5 (Romanowski et al. 2020). The probability of each overlapping was determined using the hypergeometric probability formula (Supplemental Table S1).

### Accession Numbers

LSM1a, AT1G19120; LSM1b, AT3G14080; XRN4, AT1G54490; PP2A, AT1G13320; CCA1, AT2G46830; LHY, AT1G01060; PRR9, AT2G46790; PRR7, AT5G02810; PRR5, AT5G24470; TOC1, AT5G61380; RVE8, AT3G09600; ELF3, AT2G25930.

## Data availability

Sequencing data have been uploaded to the BioProject collection and are available under accession number PRJNA775638.

## Author Contributions

D.A.C., M.J.Y. and S.P.S. conceived the project and designed the experiments; D.A.C. and S.P.S. carried out the experiments and analysed the data; R.C.M. contributed to the genome-wide analysis; D.A.C., M.J.Y. and S.P.S. wrote the article; and all authors approved the article for publication.

