## Supporting Figures for "The 5’-3’ exoribonuclease XRN4 modulates the plant circadian network in Arabidopsis"

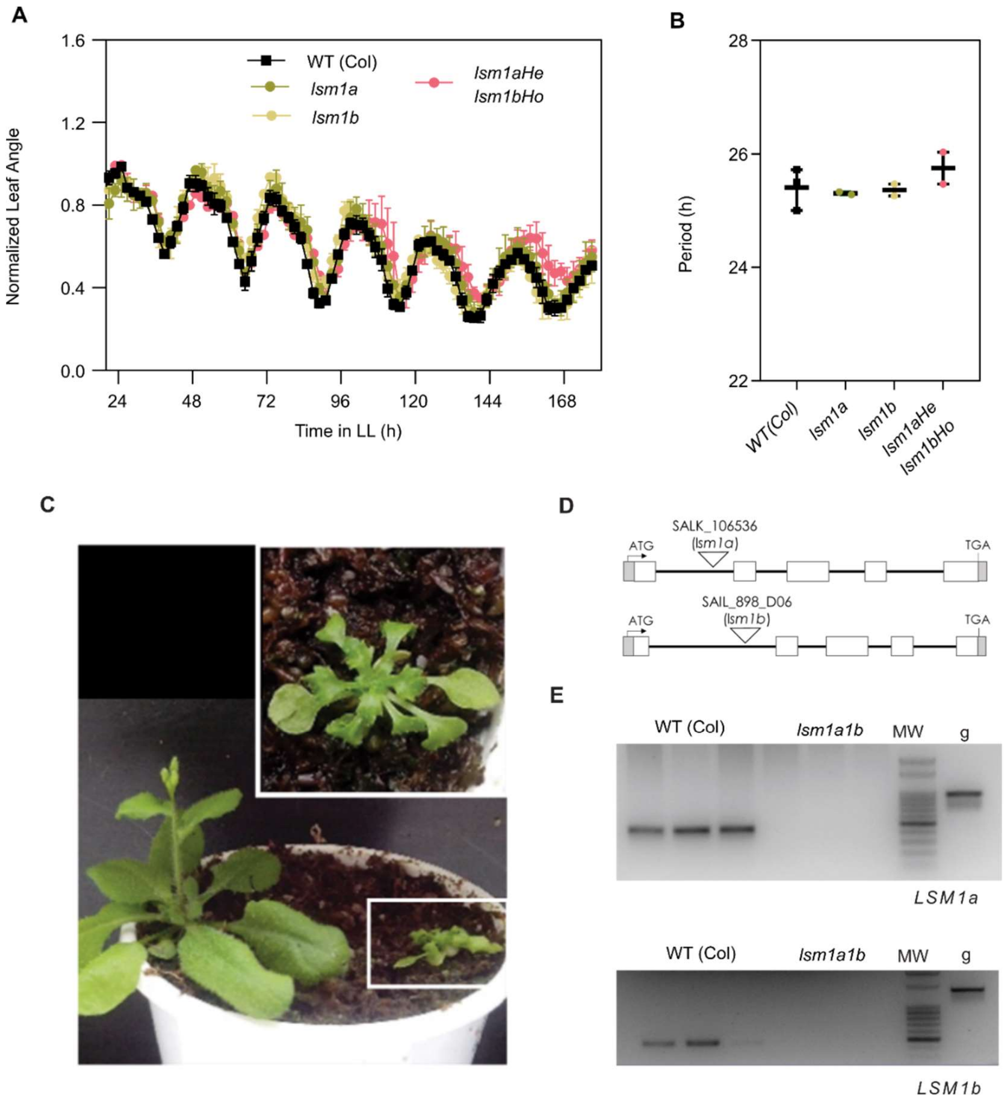


**Fig S1. Circadian phenotypes of *lsm1a* and *lsm1b* mutant plants.** (A) Leaf angles were measured for the first pair of leaves in seedlings entrained under long-day conditions (LD; 16 h light/8 h darkness), and plants were then transferred to constant light (LL) for 8 days. WT (Col), wild-type Columbia ecotype (black circles), lsm1a (olive circle), lsm1b (sand circle), lsm1aHe/lsm1bHo mutant (pink circles). (B) Period length of leaf movement rhythms estimated by fast Fourier transform–nonlinear least test (FFT-NLLS). n=2 Error bars represent mean with range. (C) Plant morphology of WT (Col) and the double lsm1a/lsm1b mutant grown in LD conditions for 30 days. The box shows an enlargement of the lsm1a/lsm1b mutant rosette phenotype. (D) Structure of the *LSM1a* and *LSM1b* gene. Introns are represented by lines and exons by rectangles; the white portions indicate coding regions and the grey, non-coding regions. The inverted triangle marks the insertion site of the T-DNA. (E) RT-PCR analysis of *LSM1a* and *LSM1b* expression indicates that *lsm1a1b* is null for both alleles. MW marker showing the 500-bp band.

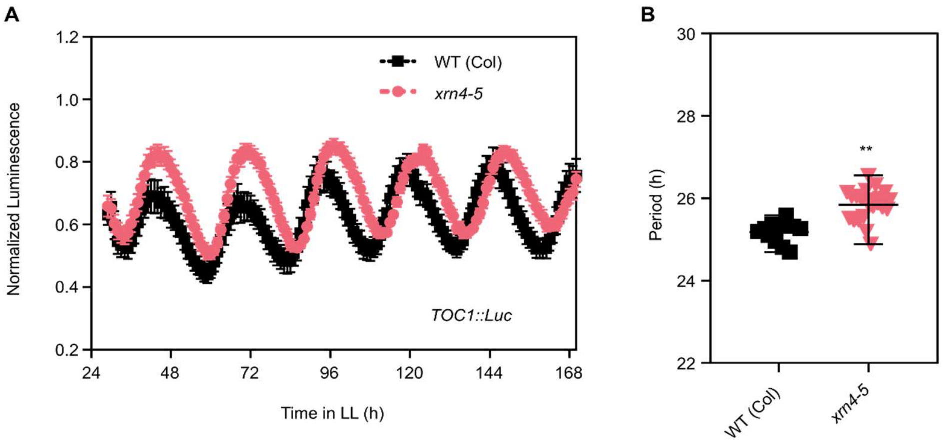


**Supplemental Figure S2. pTOC1: LUC reporter phenotype for the *xrn4-5* line.** (A) The expression of the night core clock reporter TOC1: LUC was measured for 7 d in constant light (LL) after entrainment under long-day (LD) conditions. WT (Col), wild-type Columbia ecotype (black squares, n = 11); *xrn4-5* mutant (red triangle, n = 23). (B) Bioluminescence was recorded every 2 h over the 7 days. (B) Period length of bioluminescence was estimated by fast Fourier transform– nonlinear least test (FFT-NLLS). Error bars represent mean with range. **p < 0.001 (Student t test).

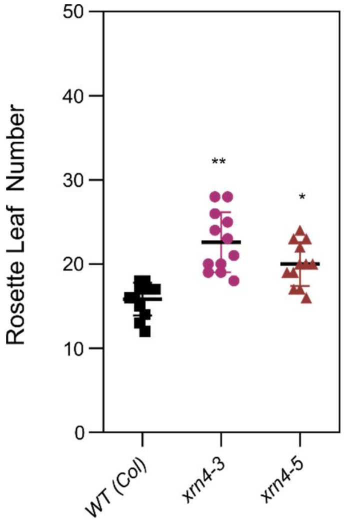


**Supplemental Figure S3. Flowering phenotype of *xrn4* mutants.** Flowering time evaluated as rosette leaf number at bolting under long-day (LD) for *xrn4-3* and *xrn4-5.* WT Col, wild-type Columbia (black squares), *xrn4-3* (purple circle), *xrn4-5* (red triangle). Error bars represent mean with range. *p<0.05 **p < 0.001 (Student t test). LD (16 h light/8 h dark).

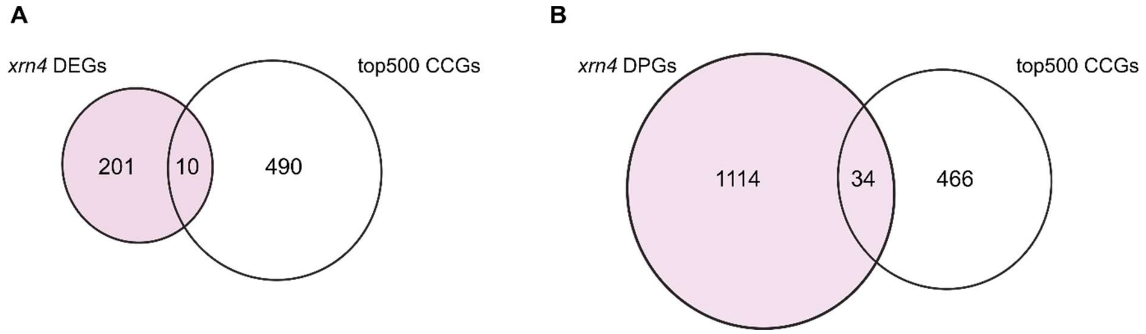


**Supplemental Figure S4. Representation of Circadian Controlled Genes (CCGs) among DEGs and DPGs.** Overlap between (A) *xrn4-3* DEGs, (B) *xrn4-3* DPGs with top500 Circadian Controlled Genes (CCGs) by amplitude. Numbers indicate the overlaps and number of DEGs, DPGs or the CCGs that passed the analysis expression filters. DEGs (Differential Expressed Genes), DPGs (Differential Post-transcriptional regulated Genes), CCGs (Circadian Controlled Genes).

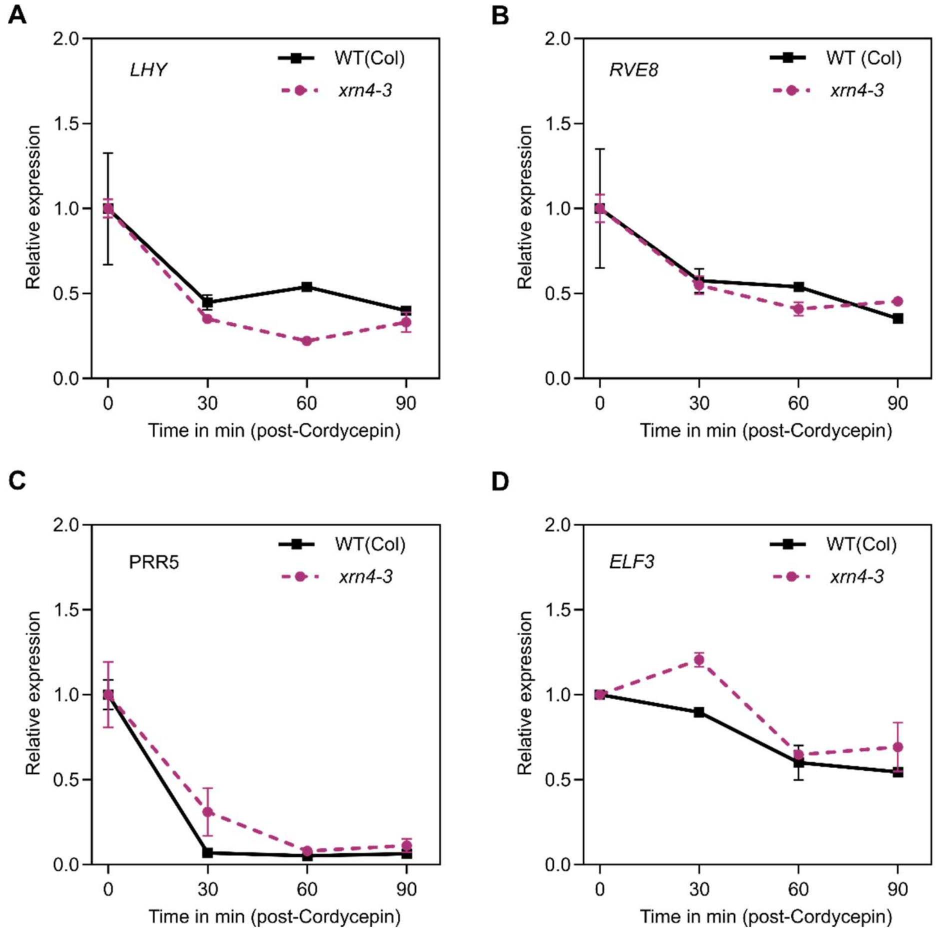


**Supplemental Figure S5. mRNA decay analysis with a cordycepin-induced transcription arrest assay.** Plant where grew for 10 days in continuous light and samples were taken every 30min from 0 to 90 min. (A) LHY, (B) RVE8, (C) PRR5, (D) ELF3. Error bars represent mean with SEM.

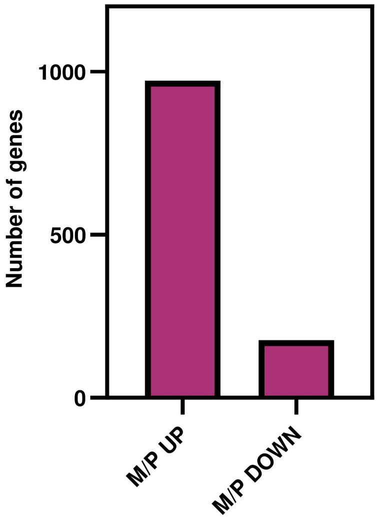


**Supplemental Figure S6.** Number of DPGs with greater or minor M/P ratio in xrn4 than WT (Col-0).

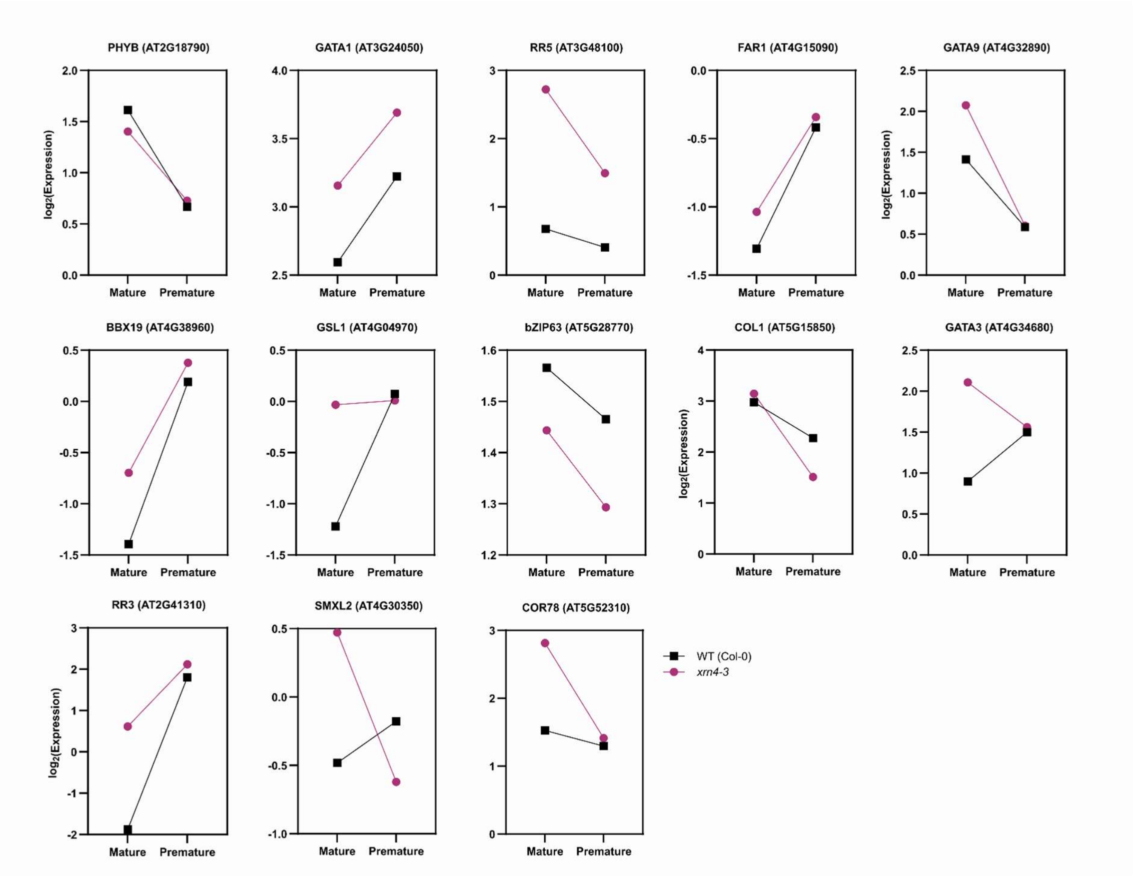


**Supplemental Figure S7. Expression of mature and premature RNA of DPGs associated with the GO term circadian rhythms in *xrn4-3*.** Log2 of RPKM of mature and premature expression estimated from exon and intron counts.
