## Supporting Tables for "The 5’-3’ exoribonuclease XRN4 modulates the plant circadian network in Arabidopsis"

**Supplemental Table S1.** Enrichment factors and p-values for overrepresentation test of the top 500 CCGs in amplitude in upregulated DEGs and DPGs. DEGs (Differential Expressed Genes), DPGs (Differential Post-transcriptional regulated Genes), CCGs (Circadian Controlled Genes).

| | **Enrichment factor** | **p-value** |
|---|---|---|
| *xrn4-3* up DEGs and CCGs | 2.2 | 0.021 |
| *xrn4-3* DPGs and CCGs | 1.2 | 0.135 |

**Supplemental Table S2.** List of primers used in this work.

| Primer | Sequence 5' -3' |
|---|---|
| LHY_Fw | GACTCAAACACTGCCCAGAAGA |
| LHY_Rv | CGTCACTCCCTGAAGGTGTATTT |
| CCA1_Fw | CATGGAAGTCTGTGTCTGACGAG |
| CCA1_Rv | TGCGGCAATACCTCTCTGGAG |
| RVE8_Fw | AAACCCTGATTTAGAATCCACTAG |
| RVE8_Rv | TGATTTGTCGCTTGTTGAGTTC |
| PRR7_Fw | GTTCCGTCAGAAGAGAAAAGAGAGG |
| PRR7_Rv | CGGCTGTTTTACGCACAAATTGG |
| PRR5_Fw | GTTAAGCCGTTGAGGAGGAATGAG |
| PRR5_Rv | GCTTTCTGCTGTCCAACACTCTC |
| TOC1_Fw | TTGGATCATCTTGCTGGGTCTCAC |
| TOC1_Rv | GCAGAGGACTCTCCGATCTTCAATC |
| ELF3_Fw | GGAAAGCCATTGCCAATCAA |
| ELF3_Rv | ATCCGGTGATGCAGCAATAAGT |
| PP2A_Fw | TAACGTGGCCAAAATGATGC |
| PP2A_Rv | GTTCTCCACAACCGCTTGGT |
